# The genetics of morphological and behavioural island traits in deer mice

**DOI:** 10.1101/443432

**Authors:** Felix Baier, Hopi E. Hoekstra

## Abstract

Animals on islands often exhibit dramatic differences in morphology and behaviour compared to mainland individuals, a phenomenon known as the “island syndrome”. These differences are thought to be adaptations to island environments, but the extent to which they have a genetic basis or instead represent plastic responses to environmental extremes is often unknown. Here, we revisit a classic case of island syndrome in deer mice (*Peromyscus maniculatus*) from British Columbia. We first show that Saturna Island mice and those from neighbouring islands are ∼35% (∼5g) heavier than mainland mice and diverged approximately 10 thousand years ago. We then established laboratory colonies and find that Saturna Island mice are heavier both because they are longer and have disproportionately more lean mass. These trait differences are maintained in second-generation captive-born mice raised in a common environment. In addition, island-mainland hybrids reveal a maternal genetic effect on body weight. Using behavioural testing in the lab, we also find that wild-caught island mice are less aggressive than mainland mice; however, lab-raised mice born to these founders do not differ in aggression. Together, our results reveal that these mice respond differently to environmental conditions on islands – evolving both heritable changes in a morphological trait and also expressing a plastic phenotypic response in a behavioural trait.

## INTRODUCTION

Islands offer some of the most tractable examples of evolutionary adaptation and diversification [1]. Because of their geographic isolation, they form self-contained environments that facilitate the interpretation of evolutionary patterns and processes. For example, many island populations of animals share consistent differences in demography, body size, reproductive rate, anti-predator behaviour, and territorial aggression, a phenomenon known as the ‘island syndrome’ [2-6]. The repeated appearance of similar traits in similar ecological conditions across taxa suggests that they evolved as adaptations during the establishment of island populations [7, 8]. Thus, one implicit assumption is that these island syndrome traits have a genetic basis, resulting from directional selection and/or founder effects during island colonization [6, 9, 10]. However, island traits could also be driven, in part or fully, by phenotypic plasticity, but the role of plasticity in island evolution has received limited attention.

A key challenge in detecting heritable components underlying island syndrome traits has been that most reports are based on experiments in the field, or especially for behavioural traits, wild-caught individuals subsequently tested in the lab (e.g. [11-14]). Detection of heritable differences in the wild, for example through the quantitative-genetic analysis of trait variance [15], can be challenging. By contrast, a genetic component can be measured if wild-derived animals are born and raised in a common environment. Using this approach, one can assess the extent to which morphological and behavioural island syndrome traits associated with islands have a heritable genetic basis.

Here, we revisit a classic case of island syndrome in deer mice (*Peromyscus maniculatus*) from western Canada. Field studies in the 1970s have shown that the ecological conditions differ for deer mice on Saturna Island off the coast of British Columbia – namely they have a higher population density, lower dispersal rate, and smaller home range size than deer mice from a nearby mainland population [16]. In addition, wild-caught island deer mice have a larger body size [16, 17] and display fewer aggressive and defensive interactions than mainland deer mice [12, 13]. Here, we investigate the extent and mechanism of genetic versus environmental components underlying morphological and behavioural island syndrome traits by measuring body size and aggressive behaviour both in wild-caught island and mainland deer mice as well as in their captive-born offspring.

## METHODS

### Body weight of museum specimens

We retrieved records (species, sex, age, body weight, body length and tail length) from museum specimens of *Peromyscus maniculatus* collected in Washington and British Columbia in the Arctos and Vertnet databases. We then filtered the dataset to ensure only adult male *P. maniculatus* were included (see Electronic Supplement Material 1 for details). Our final sample comprised 504 mice, including 53 from our own collection (see ‘Fieldwork’).

### Fieldwork

We sampled deer mice (N = 120) in the Gulf Island National Park Reserve on Saturna Island and Pender Island (*Peromyscus maniculatus saturatus*) and in the Malcolm Knapp Research Forest in Maple Ridge (*P. m. austerus* and *P. keeni*) in October 2014. Of these, we collected 101 individuals and accessioned them into the Mammal Department of the Museum of Comparative Zoology at Harvard University.

### Species assignment

Two morphologically similar species of deer mice co-occur at our mainland sampling location, *Peromyscus maniculatus* and *P. keeni.* To verify the species identity, we sequenced a partial fragment (381bp) of the *cytochrome b* gene from field-collected individuals using previously reported primers and PCR conditions [18] *P. keeni* have distinct *cytb* haplotypes from *P. maniculatus* (Suppl. Fig. 1), and these individuals were excluded from subsequent analyses.

### Admixture analysis and divergence dating

We extracted genomic DNA from 65 wild-caught mice (Saturna Island, N=28; Pender Island, N=9; mainland, N=28). We used a genotype-by-sequencing approach (ddRADseq; [19]) to obtain genome-wide markers, and then ran genetic principal component, admixture and divergence time analyses (see Electronic Supplement Material 1 for details).

### Establishment of laboratory colonies and animal husbandry

After quarantine of the wild-caught mice at Charles River Laboratories, we established colonies at Harvard University with 18 individuals from Saturna Island and 11 individuals from the Malcolm Knapp Research Forest (see Electronic Supplement Material 1 for details). Subsequent experiments were conducted using either wild-caught mice or first-, second- and third-generation captive-born mice as noted. All experiments were approved by the Harvard University Faculty of Arts and Sciences Institutional Animal Care and Use Committee (protocol #14-08-211).

### Weight measurements

Mice in the field were weighed using a Pesola spring scale to the nearest gram. Mice in the lab were weighed with a Jennings TB500 digital precision scale to the nearest 0.01g. To weigh newborn litters before their first milk meal, we isolated and monitored highly pregnant females through remote surveillance of the cage with multiple cameras. We thus were able to intercept females within 20-60 minutes of birth, and verified that pups had not yet had a meal by checking for the presence of milk in the stomach.

### EchoMRI

EchoMRI experiments were conducted in the Metabolic Core of Brigham and Women’s Hospital Boston. Frozen mice were thawed to room temperature and measured individually in a calibrated EchoMRI 3-in-1 machine.

### X-ray measurements

To measure skeletal traits, we X-ray imaged mice on a digital X-ray system in the Digital Imaging Facility of the Museum of Comparative Zoology. We used ImageJ to measure 10 skeletal elements: length of the sacrum, skull, zygomatic bone, humerus, femur, ulna, tibia, and metatarsal calcaneus as well as the width of the skull and zygomatic bone.

### F1 hybrids

We reciprocally paired one female and one male litter mate each from five island and mainland parents. Of these 10 pairs, four island female x mainland male and three mainland female x island male pairs successfully reproduced.

### Cross-fostering

Within 48 hours of birth, we exchanged pups between island and mainland parents. We fostered the same number of pups from each litter because we did not want to dilute nursing effects of island mothers, which have slightly smaller litter sizes (Suppl. Fig. 2), on offspring body weight.

### Behavioural experiments

We measured territorial aggression using a resident-intruder paradigm. To induce territoriality, residents were first paired with a female. For one set of experiments, 9 to 13 week-old residents were isolated with a female for one week. For a second set of experiments, residents were breeding males and had been paired with a female for several months. Intruders were raised in social groups with at least two other mice, and were 9 to 13 weeks old. Resident and intruder had different parents and had not previously met. We tested each resident twice against different intruders on consecutive days. We conducted all behavioural experiments during the dark phase under red light (see Electronic Supplement Material 1 for details).

To test the aggressive behaviour of males exposed to a receptive female, we used breeding males from our colony. Once a female was pregnant, we set up continuous video-monitoring of the cage, and once a litter was found, noted the approximate time of birth. Females typically reached post-partum estrus approximately 6-14 hours after the birth of the litter, as evidenced by the onset of male mating behaviour (e.g. by vigorous pursuit [20]). Males were tested approximately one hour after the onset of mating attempts and after several mating attempts had occurred.

### Behavioural analysis

We annotated videos blind to population identity using the software Observer (Noldus). We scored aggressive (lunging, pindown, upright threats, boxing, wrestling, and chasing), defensive (submission, defensive threat, and defeat), and cohesive behaviours (following, mounting, grooming, and huddling) as start-stop events performed by both the resident and the intruder.

### Statistical analysis

We fit linear models with lm {stats} or lmer {lme4} using R software [21]. *P* values were calculated using lsmeans and contrast {lsmeans} or summary {lmerTest} (see Electronic Supplement Material 1 for model details).

## RESULTS

### Island mice are heavier than mainland mice

Using data from available museum specimens, we first compared 504 weight records of adult island (N = 328) and mainland (N = 176) deer mice from 49 locations in the Strait of Georgia and coastal British Columbia. With few exceptions, island populations had a greater median weight than neighbouring mainland populations (Fig. 1A). The overall median weight of island specimens was 21g compared to a median weight of 16g in the mainland specimens (Kruskal-Wallis test, *P* < 0.0001), an increase of 31.25% (Fig. 1B). Although there was overlap in the distributions, there was only one record of a mouse more than 23g from a mainland population, whereas 30.48% of all island mice were heavier than 23g.

**Figure 1.**
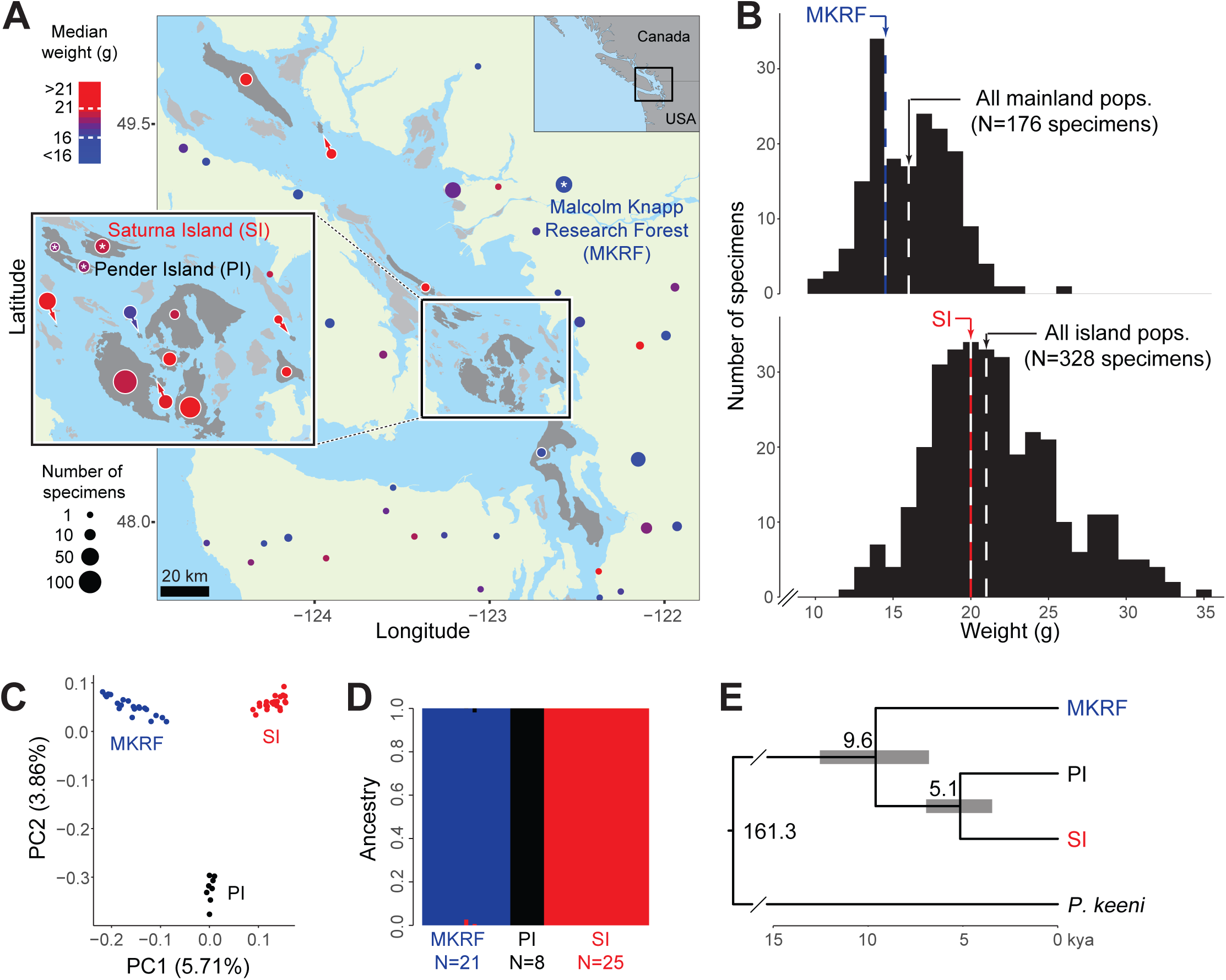
Body weight of island and mainland populations in the study area. Map of coastal British Columbia, Canada, showing the study area. The colour of collection points indicates the median weight of *Peromyscus maniculatus* populations, and the diameter of points indicates sample size. Collection points from the same island and mainland records within 8 km of each other were combined into a single point. Only adult males were included. Sampled islands are shaded dark grey, non-sampled islands light grey, and mainland areas light green. Live deer mice were collected from Saturna Island (SI) and the Malcolm Knapp Research Forest (MKRF) on the mainland to start laboratory colonies (starred). (**B**) Histograms of body weight of mainland (top) and island (bottom panel) specimens. The median weights for all populations are indicated by dashed lines. (**C**) Genetic principal component analysis of 54 deer mice from Saturna Island, Pender Island (PI) and the Malcolm Knapp Research Forest, based on 80,248 genome-wide variants. The percent of variance explained by the first and second principal component is provided. (**D**) Admixture analysis of the dataset from (C) with a cluster number of K=3. (**E**) Bayesian divergence time estimates of island and mainland *P. maniculatus* populations and the outgroup *P. keeni* based on 7,092 variants from 14 specimens. Grey bars represent 95% highest posterior density intervals. The scale bar is in thousands of years (kya).

We next focused on two populations, one island (Saturna Island) and one on the mainland (Malcolm Knapp Research Forest; Fig. 1A). We collected wild mice from these two populations (N = 21 and 23, respectively) and found that these populations largely recapitulated the general pattern of body weight observed in the Strait of Georgia. The median weight of adult Saturna Island mice was 20g (± 1 SD 16.8-23.2g) compared to a median weight of 14.5g (± 1 SD 12.7-16.3g) in Malcolm Knapp Research Forest mice (Kruskal-Wallis test, *P* < 0.0001), an increase of 37.93% (Fig. 1B).

### Island and mainland mice are genetically distinct populations

Using genetic data, we tested the genetic relatedness of our focal island and mainland populations and estimated their divergence time. We used a genotype-by-sequencing approach (ddRADseq; [19]) to obtain 80,248 genome-wide variants in a sample of deer mice from Saturna Island (N = 25), neighbouring Pender Island (N = 8), and the Malcolm Knapp Research Forest (N = 21; Fig. 1A). Using these variants, we conducted a genetic principal component analysis (PCA) and found that these populations are genetically separable (Fig. 1C). In addition, none of the individuals examined showed evidence of recent admixture, suggesting that gene flow between island and mainland populations, as well as between neighbouring islands, is limited (Fig. 1D). Lastly, we applied Bayesian divergence time estimations to a subset of high coverage genetic variants (N = 7,092) and specimens (N = 14) to obtain a time-calibrated phylogenetic tree of these populations, using *Peromyscus keeni* as an outgroup (Fig. 1E). Using this approach, we estimated island mice diverged from mainland mice approximately 9.6 (6.8-12.5, 95% highest posterior density interval) thousand years ago (kya), while Pender Island and Saturna Island mice diverged approximately 5.1 (3.5-6.9) kya. These data are consistent with a postglacial isolation of deer mice on islands in the Strait of Georgia [22] and little to no ongoing gene flow between island and mainland or among island populations.

### Differences in body weight appear heritable and are driven by both differences in body length and proportional lean mass

To test whether differences in weight between Saturna Island and mainland mice observed in the field were retained across generations in a common environment, we established laboratory colonies of these two populations and then compared a set of wild-caught (N = 21) and captive-born (N = 25) mice. We used captive-born mice that were themselves born to captive-born parents (as two generations in captivity should minimize environmental, maternal and other non-heritable variation), and focused our analyses on male mice because the difference in weight was greater in males than in females (Suppl. Fig. 3). We found that the magnitude of difference in median weight between island and mainland mice observed in the field was retained in laboratory mice (island mice are 37.93% bigger in the field and 34.02% bigger in the lab; Fig. 2A); however, the overall weight of the mice was higher in the laboratory (captive-born mice were on average 34.99% heavier than wild-caught mice; Fig. 2A), likely due to an enriched and *ad libitum* diet.

**Figure 2.**
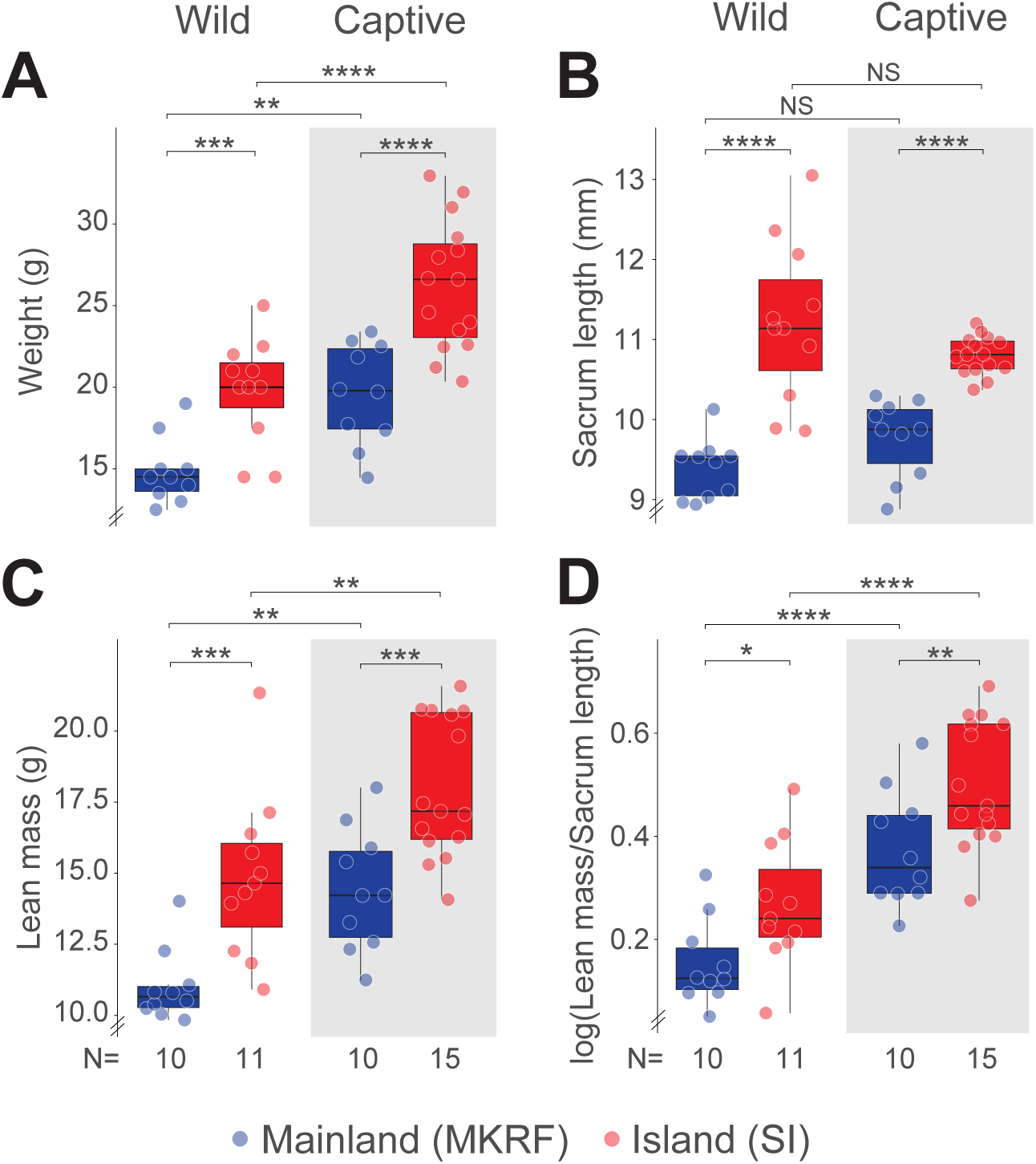
The genetic basis of body size traits in island mice. Body weight, (**B**) sacrum length, (**C**) lean mass, and (**D**) size-corrected lean mass in male wild- and captive-born island and mainland mice. See Methods for statistical details. Statistical significance evaluated by linear fixed effects models. For all figures, box plots indicate median, interquartile range (IQR), and at most ± 1.5 IQR from hinges. NS=not significant, * *P* < 0.05, ** *P* < 0.01, *** *P* < 0.001, **** *P* < 0.0001.

We next asked how island mice attain their larger body weight and hypothesized that they might be heavier both because they are larger (i.e. longer) and/or because they are heavier relative to their body length. To distinguish among these possibilities, we first X-rayed mice to measure a set of 10 skeletal traits: length of the sacrum, skull, zygomatic bone, humerus, femur, ulna, tibia, and metatarsal calcaneus as well as the width of the skull and zygomatic bone. We found that median sacrum length, a proxy for body length, was approximately 14.39% larger in island than mainland mice for both field and lab individuals, but did not differ between field and lab populations (Fig. 2B), suggesting that island mice are longer than mainland mice but that environmental conditions (i.e. field versus lab) have little effect on body length. This pattern was largely recapitulated in other skeletal traits: size-corrected proxies for head length/width and several limb bones showed that island mice have disproportionately smaller heads and shorter front and hind legs in both field and lab populations, and little difference between field and lab conditions (Suppl. Fig. 4).

We next quantified body composition in field and laboratory mice using EchoMRI analyses. We found that island mice had more lean mass (Fig. 2C), but not fat mass (Suppl. Fig. 5), than mainland mice. Moreover, this increase in lean mass was not simply due to differences in body length (Fig. 2D). In addition, lean mass and relative lean mass was greater in captive-born than in wild-caught mice from the same location (Fig. 2C, D), suggesting that these traits are influenced by the environment. Collectively, these data show that island mice are heavier both because they are larger (i.e. longer) and because they have disproportionately more lean mass than mainland mice. In addition, because these trait differences are maintained both in a common environment and across generations, they likely have a strong genetic basis.

### Island mice are born heavier with growth pulses at birth and weaning

To determine when these weight differences emerge across ontogeny, we weighed litters within 24 hours of birth, and for a subset continued weighing them every 4 days until weaning. We noted that island mice have slightly smaller litter sizes than mainland mice (Suppl. Fig. 2; Kruskal-Wallis test, *P* = 0.034); as litter size is often inversely correlated with birth weight, we thus included litter size as an explanatory variable in subsequent analyses. Because individual pups are difficult to track over time, we focused on comparisons among litters. We first found that island litters are heavier than mainland litters already at birth (Fig. 3; linear fixed effects model, t = −6.259, *P* = <0.0001). This difference is not simply due to island mice ingesting more milk because island litters were already heavier before their first milk meal (Suppl. Fig. 6; repeated measures linear mixed effects model; t = 6.901, *P* = 6.96e-5 [pre vs. post-feeding]; t = 3.602, *P* = 0.004 [Strain]). Furthermore, growth at birth in island mice is greater than in mainland mice (Suppl. Fig. 7A; linear fixed effects model, t = −2.852, *P* = 0.033), but this significant difference disappears by postnatal day 4 (P4), and only reappears around weaning at P23 (Suppl. Fig. 7A; linear fixed effects model, t = −2.748, *P* = 0.039). Notably, in both strains the post-natal growth peak at P4 slows down by P12, and growth only resumes and exceeds earlier levels subsequent to P16. Together, these results suggest that growth pulses around birth, and with the onset of sexual maturity at weaning, underlie the differences in weight between island and mainland mice.

**Figure 3.**
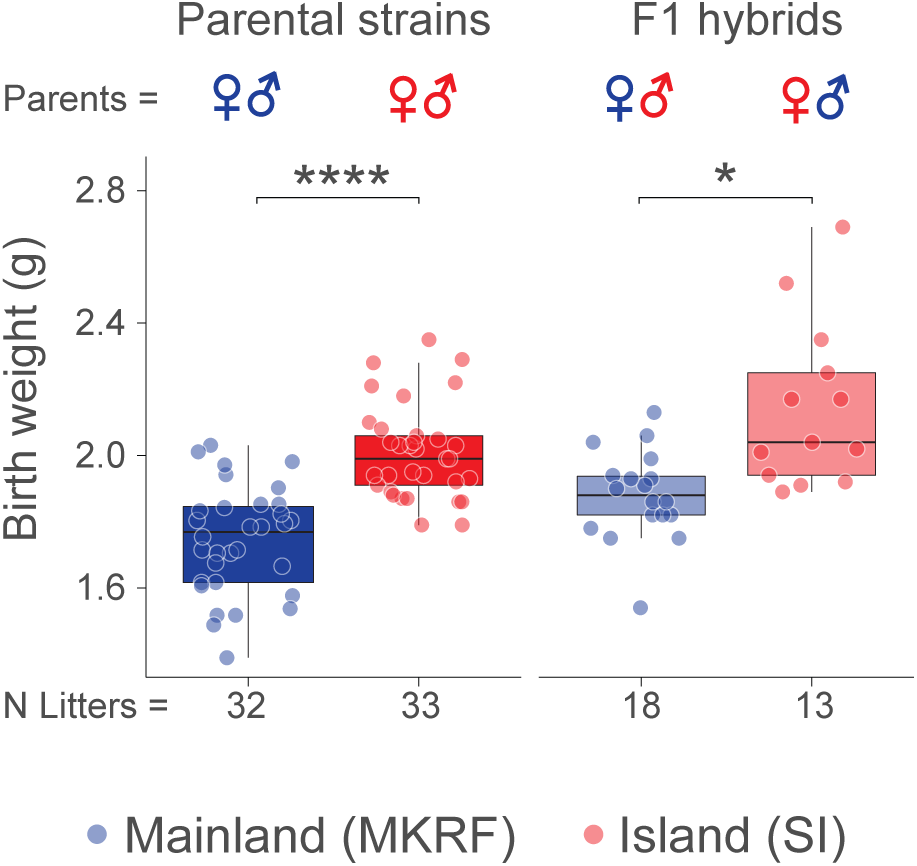
A maternal effect contributes to birth weight in island mice. Birth weight in island and mainland litters (left panel) and F1 hybrids by maternal strain (right panel). Points represent the mean weight of litter. Statistical significance evaluated by linear fixed effects model with litter size as an explanatory variable (see Methods for details). * *P* < 0.05, **** *P* < 0.0001.

### A maternal effect contributes to birth weight and early growth in island mice

We next asked if these differences in weight are influenced by a maternal genetic effect by comparing first-generation (F1) hybrid litters born to mothers from either island or mainland populations. We first reciprocally paired litter mates from representative island and mainland parents. As before, we weighed litters at birth and then a subset every 4 days until weaning. We found that F1 litters born to island mothers are on average 14.11% heavier at birth than F1 litters born to mainland mothers (Fig. 3; linear fixed effects model with litter size as an explanatory variable, t = −2.386, *P* = 0.0241), even when we controlled for family effects (wild cluster bootstrapped generalized linear model, 95% CI = 0.0022, 0.1749, *P* = 0.029). Furthermore, growth in F1 hybrid litters born to island mothers was significantly greater at birth (Suppl. Fig. 7B; linear fixed effects model, t = −4.114, *P* = 0.0005) and at P4 (Suppl. Fig. 7B; linear fixed effects model, t = −3.535, *P* = 0.0034), but we only saw a trend at P4 when we controlled for family effects (wild cluster bootstrapped generalized linear model, 95% CI = −0.1076, 0.6122, *P* = 0.069). Growth was not different on days 8-20 or at weaning (Suppl. Fig. 7B; linear fixed effects model, t = 1.085, *P* = 0.84). Finally, in F1 adults in which sex could be determined, we found a larger difference between offspring with island versus mainland mothers in male compared to female hybrids (Suppl. Fig. 7C, D). Together, these results suggest that the growth difference in island versus mainland mice around birth, but not at the onset of sexual maturity at weaning, is likely governed by a maternal effect.

### Parental care does not affect early growth differences

We next tested if differences in parental care contribute to the differences in growth observed between island and mainland mice. To address this hypothesis, we conducted a cross-fostering experiment by swapping island and mainland litters (i.e. island pups raised by mainland parents and vice versa) within 48 hours of birth. We then weighed the pups every 4 days until weaning (P23). We found little evidence that cross-fostering affected growth in either island or mainland mice (Suppl. Fig. 8A, B). For example, mainland litters raised by island parents grew the same (days 0 to 20) or slower (day 23) compared to non-fostered mainland litters (Suppl. Fig. 8A). These data suggest that the increases in growth in island litters and F1 hybrids born to island mothers are not due to increased parental care by island parents.

### Wild-caught island mice are less aggressive than wild-caught mainland mice

We next compared levels of territorial aggression in both wild-caught and captive-born mice from island and mainland populations. We first developed a behavioural assay in the laboratory to test aggression levels in male mice based on the resident-intruder paradigm (Fig. 4A). Briefly, males were co-housed with females to initiate territoriality. Females were removed immediately before the trial from the cage. A conspecific, socially-housed male intruder was then introduced into a closed-off section of the cage. After habituation, the resident and intruder were allowed to interact, and behaviour was video-recorded for 15 minutes. We conducted a second trial against a different intruder one day later. We then quantified a suite of social behaviours [23, 24] as state (duration) events from the videos, and averaged resident behaviours across trials.

**Figure 4.**
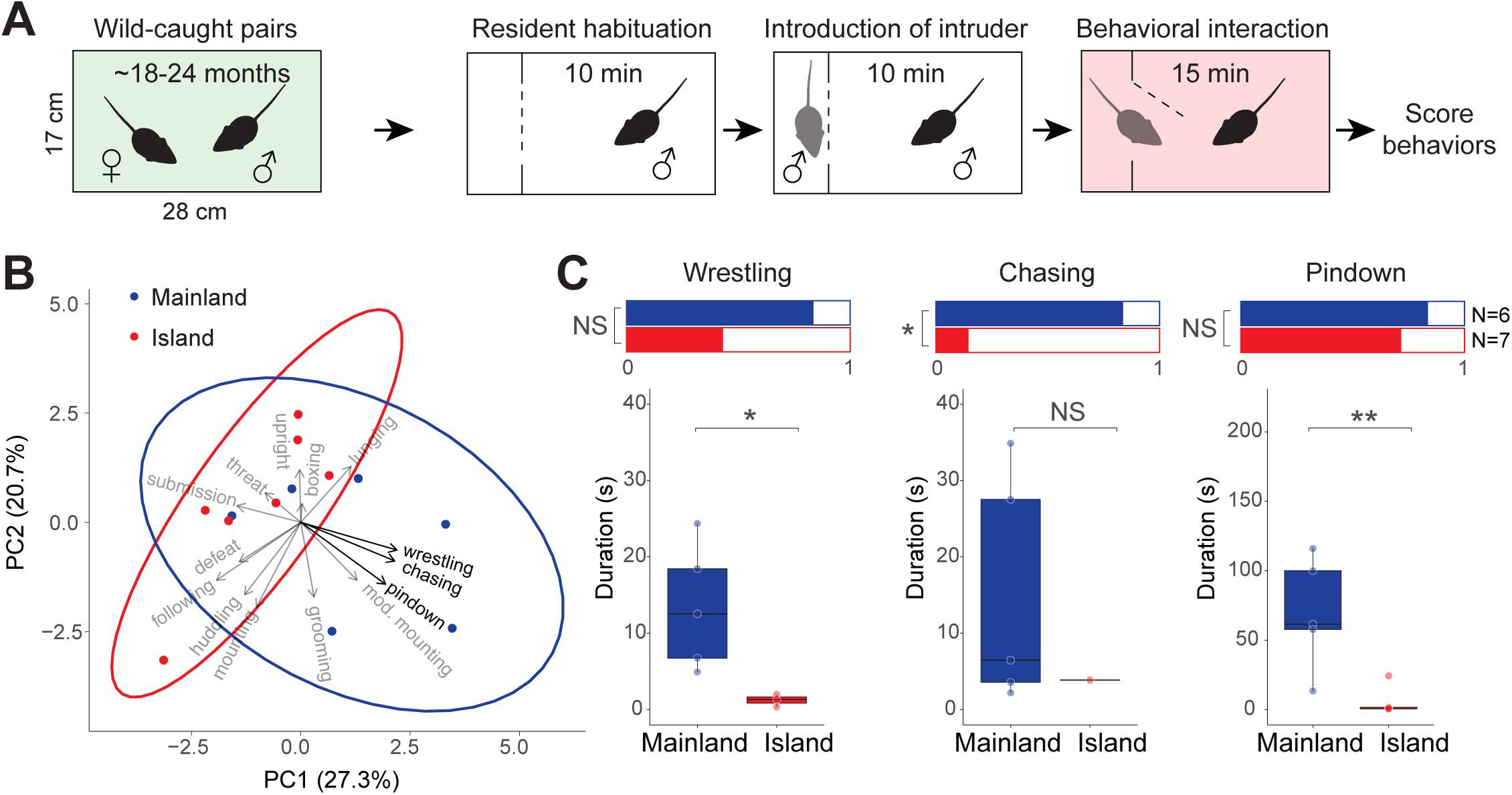
Territorial aggression in wild-caught island and mainland mice. (**A**) Schematic of the behavioural assay. For each resident, we ran the assay twice separated by a day using two different conspecific intruders. We scored aggressive, defensive, and cohesive behaviours and averaged scores across trials. (**B**) Principal component analysis (PCA) of the duration of wild-caught resident behaviours. Ellipses represent 95% confidence intervals. Arrow directions indicate how behaviours contribute to the two PCs, and arrow lengths are proportional to the strength of contribution. Behaviours highlighted in bold were selected for additional analysis. (**C**) Aggressive behaviour in wild-caught founder animals. Bar graphs (top) show the proportions of animals engaging in wrestling, chasing, and pindown behaviours. Boxplots (bottom) show duration of each behaviour for those individuals that engaged in that behaviour. Sample sizes are provided. Statistical significance of proportions and durations evaluated by generalized linear models and linear fixed effects models, respectively (see Methods for details). NS=not significant, * *P* < 0.05, ** *P* < 0.01.

We first tested the male mice that were collected in the wild and paired to establish laboratory colonies (N=7 island, N=6 mainland). At the time of behavioural testing, these individuals had been co-housed for ∼1.5-2 years with a female in the lab. We performed a principal component analysis (PCA) to reduce the dimensionality of the behavioural dataset (Fig. 4B). The first two principal components (PCs) explained a cumulative proportion of 47.92% of the variance; 95% confidence intervals for the two strains only partially overlapped, and 67% of mainland animals were outside the confidence interval for island mice. The behaviours that contributed most to the first PC were wrestling duration (12.94%), chasing duration (12.65%), and pindown duration (11.42%).

Using this dataset, we tested for differences in these behaviours between wild-caught island and mainland mice. We used a two-stage hurdle model approach to separately test for differences in the proportion of animals displaying a behaviour as well as the number and duration of bouts once a behaviour was initiated. We found no significant difference in the proportion of wild-caught island and mainland mice showing wrestling behaviour (83.3% vs. 42.86%, logistic regression model, z = −1.421, *P* = 0.155), but mainland animals that wrestled did so significantly longer than island animals (Fig. 4C, Suppl. Fig. 9A; linear fixed effects model, t = −2.512, *P* = 0.0458). For chasing, significantly more mainland animals chased than island animals (83.3% vs. 14.29%, logistic regression model, z = −2.211, *P* = 0.027), but we found no difference in duration once animals chased (Fig. 4C, Suppl. Fig. 9A; linear fixed effects model, t = −0.666, *P* = 0.5418). Similar to wrestling behaviour, we found no significant difference between populations in the proportion of animals exhibiting pindown behaviour (83.3% vs. 71.43%, logistic regression model, z = −0.503, *P* = 0.615), but mainland animals that exhibited pindown behaviour did so significantly longer than island animals (Fig. 4C, Suppl. Fig. 9A; linear fixed effects model, t = −3.460, *P* = 0.0086). These differences in aggression could not be explained by differences in the time that animals had spent in captivity, the time since a litter was last sired, or the weight difference between resident and intruder (Suppl. Fig. 10). Together, these results suggest that wild-caught island mice show reduced territorial aggression compared to wild-caught mainland mice, consistent with previous studies on wild mice from these populations [12, 13].

### Captive-born mice tested in standard conditions are not aggressive

To determine if the behavioural differences observed in wild mice were retained in a controlled laboratory environment and over generations, we next compared aggression in captive-born island and mainland mice. We initially paired residents with a virgin female for one week prior to the behavioural testing (Fig. 5A), a well-established procedure to induce territoriality and aggression in laboratory mice [25]. However, both captive-born island and mainland mice displayed low levels of aggression in this assay (Fig. 5B, Suppl. Fig. 9B). We found neither a significant difference in the proportion of island versus mainland mice that displayed wrestling (21.73% vs. 24.39%, logistic regression model, z = 0.240, *P* = 0.81), chasing (30.43% vs. 17.07%, logistic regression model, z = − 1.227, *P* = 0.22), or pindown (47.83% vs. 53.66%, logistic regression model, z = 0.448, *P* = 0.654), nor did we find a significant difference in the duration of wrestling (linear fixed effects model, t = −1.938, *P* = 0.0746), chasing (linear fixed effects model, t = −1.432, *P* = 0.1777), or pindown (linear fixed effects model, t = 0.268, *P* = 0.7905) once animals engaged in these behaviours. These results are in stark contrast to differences observed in wild-caught mice; both island and mainland lab-reared animals behaved similarly to the less aggressive wild-caught island mice.

**Figure 5.**
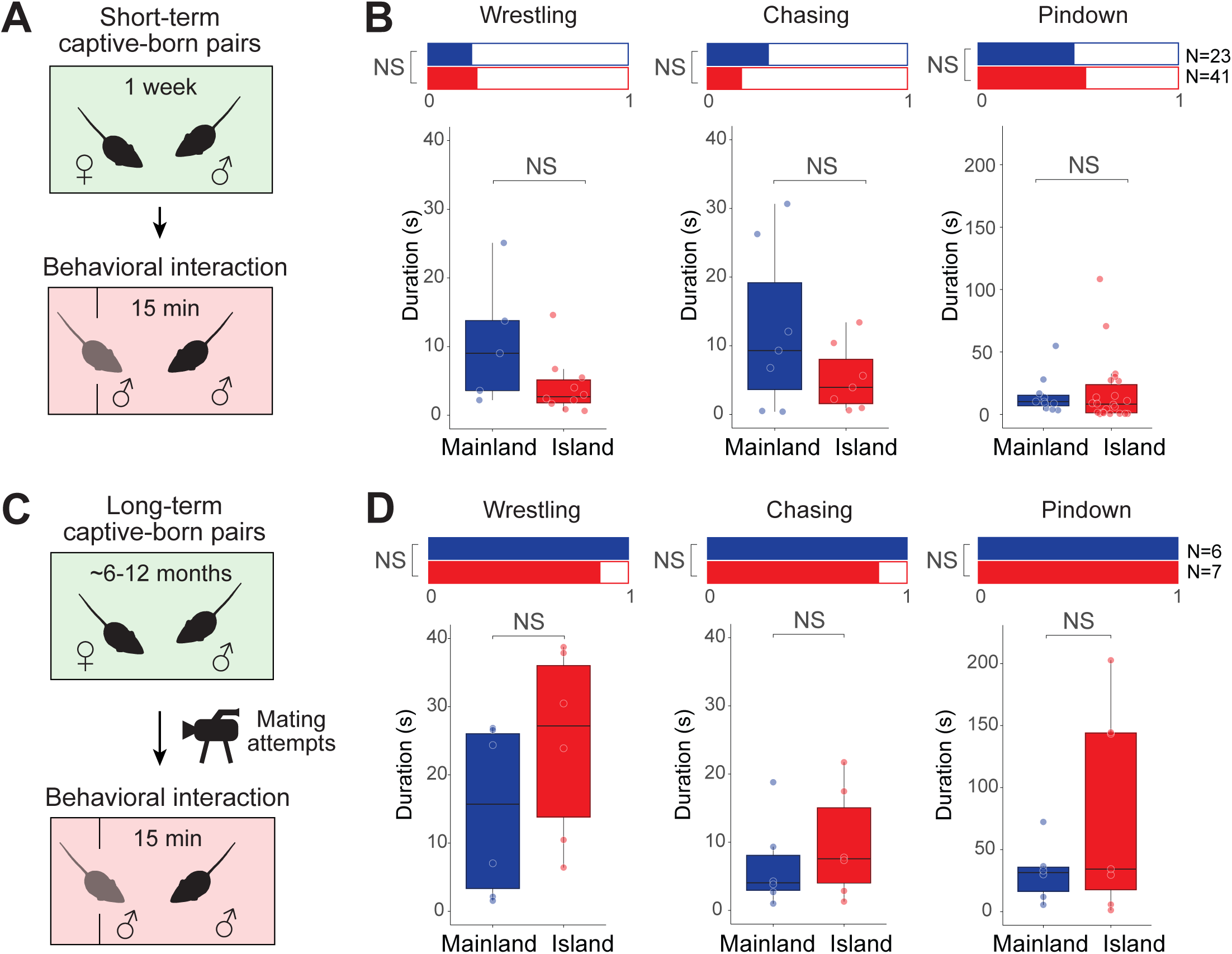
Territorial aggression in captive-born island and mainland mice. (**A**) Schematic of the resident-intruder behavioural assay, a standard paradigm to induce aggression in laboratory mice, using captive-born males paired with a female for one week. For each resident, we ran the assay twice separated by a day using two different conspecific intruders. (**B**) Aggressive behaviour in short-term pairings. Bar graphs (top) show the proportions of animals engaging in wrestling, chasing, and pindown behaviours. Boxplots (bottom) show duration of each behaviour for those individuals that engaged in that behaviour. Statistical significance of proportions and durations evaluated by generalized linear models and linear fixed effects models, respectively (see Methods for details). (**C**) Schematic of the behavioural assay using captive-born males paired with a female for 6-12 months, and then testing males approximately one hour after the onset of mating attempts and again one day later. (**D**) Aggressive behaviours in long-term pairings. Bar graphs (top) show the proportions of animals engaging in wrestling, chasing, and pindown behaviours. Boxplots (bottom) show duration of each behaviour for those individuals that engaged in that behaviour. Statistical significance of proportions and durations evaluated by generalized linear models and linear fixed effects models, respectively (see Methods for details). Sample sizes are provided. NS=not significant.

### Reproductively active captive-born mice do not differ in aggression levels

While one week of pairing with a female typically induces sexual activity and territoriality in male laboratory mice, it may not be sufficient for deer mice. To explore this possibility, we tested for a subset of residents when they sired their first litter. We found that the majority of males sired litters approximately three weeks after they were paired (Suppl. Fig. 11A), suggesting that few mice had become sexually active by the time of testing after only a single week. Furthermore, we identified mice among this subset that had not sired a litter by the time of first testing, and re-tested them after they had sired and been co-housed with a litter until weaning. As expected, residents wrestled significantly more after they had reproduced (Suppl. Fig. 11B; repeated measures linear mixed effects model; t = 2.578, *P* = 0.0327 [Trial]; t = 0.337, *P* = 0.7462 [Strain]), suggesting that reproductive experience is crucial to aggression in these deer mouse populations.

As aggression in male deer mice peaks at the time of and just following copulation with a receptive female [26-28], we next aimed to test for island-mainland differences in aggression at this time. We used first-generation captive-born breeding animals that had been paired for 6-12 months and took advantage of the fact that females undergo post-partum oestrus (Fig. 5C). We identified pregnant females in our colony and continuously video-monitored their behaviour in their home cage. We found that male mice will begin to attempt copulation about 12 hours after the birth of a litter for about 1-2 hours (Suppl. Fig. 12). Therefore, we tested male mice in our behavioural assay approximately one hour after the onset of copulation and again one day later.

As expected, aggression levels were very high in reproductively active mice (Fig. 5D, Suppl. Fig. 9C). Nearly all residents of both populations exhibited wrestling, chasing, and pindown behaviours. Surprisingly, however, there was no significant difference in the proportion of island and mainland mice exhibiting wrestling (100% vs. 85.71%, logistic regression model, z = −0.003, *P* = 0.998), chasing (100% vs. 85.71%, logistic regression model, z = −0.003, *P* = 0.998), or pindown behaviour (100% vs. 100%, logistic regression model, z = 0, *P* = 1), nor was there a difference in the duration of wrestling (linear fixed effects model, t = 1.309, *P* = 0.2199), chasing (linear fixed effects model, t = 0.725, *P* = 0.4851), or pindowns (linear fixed effects model, t = 1.409, *P* = 0.186). Thus, although we were able to induce higher aggression levels in captive-born mice, we still found no behavioural differences between captive-born island and mainland mice, implying that the aggression differences observed in wild-caught mice are driven, at least in large part, by environmental effects.

## DISCUSSION

Here, we investigated the genetic basis of morphological and behavioural traits associated with the island syndrome in deer mice from British Columbia. Previous work in the 1970s reported that wild-caught Saturna Island deer mice had higher body weight [16, 17] and reduced aggression levels compared to mainland mice [12, 13]. We recapitulated these differences in our populations of wild-caught deer mice. First, we found that body weight is indeed higher in Saturna Island (as well as most other islands in the region) than mainland mice, and this difference is driven by both the increased body length and disproportionately larger lean mass of Saturna Island mice. Second, we also found striking differences in aggression: wild-caught island mice showed reduced aggression in several behaviours compared to mainland mice. Thus, Saturna Island mice in the wild indeed conform to the predicted island syndrome phenotype. However, because measurements were taken using wild-caught individuals, it was unclear if these morphological and behavioural differences had a genetic basis or instead were plastic responses to environmental extremes [12, 13].

We thus established wild-derived colonies in the laboratory to minimize environmental variation on these traits. Using these colonies, we found that body size differences persisted in captive-born mice raised in common conditions, suggesting these differences were not driven by environmental factors and instead appeared largely heritable. Although examples of density-dependent plasticity in body weight are known from natural populations (e.g. [29]), our results are consistent with other studies that have demonstrated the genetic basis of gigantism and skeletal evolution in island rodents [30-32]. In addition, our data on island-mainland hybrid growth showed that the larger body weight in island mice was also partly driven by a maternal genetic effect. The maternal effect was not recapitulated by cross-fostering, suggesting that it cannot be explained simply by post-natal nursing differences. A study in island and mainland deer mice from California showed that four day old mainland embryos that were transferred into island foster mothers had a significantly increased neonatal weight [31], suggesting that increased maternal uterine investment also contributes to the large size of Saturna Island deer mice. However, we cannot exclude the possibility of maternal genomic imprinting (e.g. [33]) without an embryo transfer experiment. Regardless, it is clear that both offspring and maternal genotypes driving larger body size have evolved in island deer mice.

Unlike body size differences, however, our laboratory-based behavioural experiments demonstrated that aggression levels in captive-born island and mainland deer mice remained indistinguishable across a spectrum of low to high overall aggression intensity. These results suggest that additional environmental factors are necessary to induce the aggression differences observed in wild-caught mice reported previously [12, 13] and replicated in our study. One important factor may be population density [5, 6, 34]. Consistent with our findings in *Peromyscus maniculatus*, increased body size and reduced aggression have been documented in other island rodents with high population density (e.g. *Microtus breweri* [35, 36] and *Myodes glareolus skomerensis* [37]). Population density in island rodents is typically increased, likely because of the absence of emigration, fewer competing species and thus more available resources, and fewer predators on islands. The rate of social interactions, and thereby also the cost of territorial defence, likely increase with density. In addition, reproductive output in rodents is often adjusted to population density. Rodents in high density island populations could thus face pressure to reduce direct conflict in favour of indirect competition (territorial defence hypothesis) and to reallocate resources to produce fewer, but more competitive offspring (reallocation hypothesis).

The evolution of these island syndrome traits can be framed in the context of phenotypic plasticity (e.g. [38-41]). Plastic phenotypic changes are often considered primarily during the initial stages of the colonization of novel habitat. Phenotypic plasticity can transiently reduce the intensity of selection and create a time lag between the initial exposure to novel environments and the necessity for genetic adjustment [15, 42]. This has also been noted in the literature on the island syndrome; for example, it has been suggested that “environmental variation [on islands] may cause ‘purely phenotypic’ changes within a single generation [short-term changes] and genetically determined changes over evolutionary time [long-term changes]” [6]. Because we observed a capacity for behavioural plasticity in the extant population on Saturna Island, we speculate that plasticity in territorial behaviour may have existed in ancestral deer mice and may have been critical to colonize Saturna Island and sustain populations at high density. This is consistent with growing evidence that phenotypic plasticity especially in behaviour can promote population persistence and adaptation in these initial stages of colonization, because behavioural traits are thought to be more sensitive to environmental factors than morphology and may thus respond first in new environments [41, 43-45].

Over evolutionary time, it is thought that selection would then act on heritable differences in island traits [46]. Eventually, through a process of genetic assimilation, a trait would become genetically and developmentally canalized, and environmental plasticity would be lost [47]. Studies that have demonstrated phenotypic plasticity in island traits and genetic change over evolutionary time have focused primarily on morphological traits [48-50]. Consistent with these reports, the relative body size difference between Saturna Island and mainland mice was retained in captivity and thus similarly may have undergone genetic assimilation. This raises the interesting possibility that behavioural plasticity in these island mice preceded morphological evolution, and that this initial behavioural plasticity enabled the selective environment, that is, high population density, in which parental and offspring genotypes driving larger offspring body size became advantageous, lending further support to the idea that behavioural plasticity can affect the evolution of other traits, especially in morphology [45, 51-53].

Unlike the body size differences, aggression levels in island deer mice continue to be primarily driven by environmental parameters such as population density, raising the question of why plastic behaviours do not necessarily undergo genetic assimilation themselves [40, 43-45]. One may argue that highly plastic traits inherently hinder their own evolution by reducing the intensity of selection on underlying genetic variation [54]. But while plasticity may slow down genetic evolution initially, selection is expected to act on genetic variation eventually, leading to genetic accommodation of plastic behavioural phenotypes.

Adaptive evolution is the result of selection on heritable variation, and both the strength of selection and the extent of heritability can influence trait evolution. One explanation could thus be that behaviour in general is simply less heritable than morphology. However, data on the heritability of behaviour and morphology do not unanimously support this hypothesis [55, 56]. Rather, heritability of different behaviours may vary substantially. For example, similar to our findings, a comparative study of both captive and wild insular macropodid marsupials suggested that loss of some anti-predator behaviours may be heritable, while loss of others is more phenotypically plastic [57]. Moreover, selection on certain behaviours, like aggression, may be labile over time (e.g. if selection was density-dependent and density changed periodically). Thus, if selection on aggression was weak and/or heritability low, 10 thousand years (or ∼25 thousand generations) simply may not be enough time for a behavioural difference to fix between island and mainland populations.

In summary, while the island syndrome has been reported in a wide range of organisms, few studies have tested if these traits are genetically encoded. More research is needed into the drivers, sequence, and mechanisms of trait evolution during the colonization of islands and subsequent establishment of populations to fully understand one of the most iconic and widespread patterns of repeated evolution in nature.

## DATA ACCESSIBILITY

Raw sequence data (fastq files) and mtDNA sequences were submitted to the Sequence Read Archive (PRJNA508981) and GenBank (MK265122-MK265240), respectively. Morphological and behavioural data will be deposited into the Dryad Digital Repository. Museum specimens were accessioned into the Harvard Museum of Comparative Zoology Mammal Department (69713-69813).

## AUTHORS’ CONTRIBUTIONS

FB and HEH conceived the study, FB carried out the experiments and analysed the data supervised by HEH, and FB and HEH wrote the paper.

## COMPETING INTERESTS

We declare no competing interests.

### ACKNOWLEDGEMENTS

We thank J. Chupasko for help planning field work and permits; N. Bedford, E. Hager, and K. Turner for assistance in the field; and J. Kenagy for help with field logistics. We also thank H. Wellington for generating the X-ray images and taking skeletal trait measurements; A. Shultz for advice on the population-genetic analysis; M. Omura for help accessioning specimens; and A. Nelson for advice on the behavioural assay design. K. Pritchett-Corning and the Office of Animal Resources at Harvard University were instrumental in setting up breeding colonies from wild-caught mice. Data science specialists S. Worthington and S. Goshev at the Institute for Quantitative Social Science, Harvard University, provided statistical support. Fieldwork was conducted with permits from the Gulf Island National Park Reserve (GINP-2014-17276), the Pender Island Parks and Recreation Commission, the Malcolm Knapp Research Forest at the University of British Columbia, and the Ministry of Forests, Lands and Natural Resource Operations (NASU14-154228, NASU14-154230). Fieldwork was supported by a Putnam Expedition Grant of the Harvard Museum of Comparative Zoology. FB was supported by a HHMI International Student Research Fellowship, a Joan Brockman Williamson Graduate Research Fellowship, and a Herchel Smith Graduate Fellowship. HEH is an Investigator of the Howard Hughes Medical Institute.

## Electronic Supplement Material 1

### Body weight of museum specimens

To minimize confounding effects of reproductive status and age, we excluded females and specimens of unknown sex, juveniles, subadults, and embryos. We then removed specimens with a body length ≤ 70 mm and/or a tail length ≤60 mm, which represents minimum values in a sample of adult *P. maniculatus* we collected in British Columbia (see below). To remove specimens with out-dated taxonomy that no longer are classified as *P. maniculatus*, notably *P. keeni* [1], we filtered out records labelled as *P. keeni* or any of its synonyms or subspecies. We also excluded specimens with both a tail/body ratio of ≥ 1.1 and a tail length ≥ 95mm, which are morphological criteria to identify *P. keeni* [2].

### Admixture analysis and divergence dating

Genomic DNA was extracted using an AutoGenPrep 965 (AutoGen), and the quality of extractions was verified with Quant-iT (Thermo Fisher Scientific). We digested genomic DNA samples with NlaIII and MluCI enzymes (New England Biolabs) and ligated fragments to biotinylated barcoded adapters. We size-selected for 216-276bp fragments using a PippinPrep (Sage Science), cleaned fragments with Streptavidin-coupled M-280 Dynabeads (Thermo Fisher Scientific), and PCR amplified fragments using 12 uniquely indexed PCR primers with Phusion DNA polymerase (New England Biolabs). After a bead clean-up and evaluation of the library quality on a 2200 TapeStation (Agilent Technologies), we generated 125bp paired-end reads with a standard v4 run on an Illumina HiSeq 2500 sequencer.

We used a custom-made pipeline to demultiplex reads and combine them by sample into one R1 and R2 file. We then used BWA 0.7.15 [3] to map reads to an in-house *de novo* assembly (v2.1) of the *P. maniculatus bairdii* (strain BW) reference genome. After calling variants for each sample separately with GATK 3.5 [4] using “HaplotypeCaller” (including options -minPruning 1, -minDanglingBranchLength 1, and -het 0.005), we produced one vcf file containing all samples with GATK’s “GenotypeGVCFs” (including option het −0.005). After quality filtering (QualByDepth (QD) < 2; FisherStrand (FS) > 60 for SNPs, > 200 for indels; RMSMappingQuality (MQ) < 40; MappingQualityRankSumTest (MQRankSum) < 12.5; ReadPosRankSumTest (ReadPosRankSumTest) < -8 for SNPs, < -20 for indels; StrandOddsRatio (SOR) > 3 for SNPs, > 10 for indels), we used vcftools [5] to explore depth and missingness. We then removed samples of *P. keeni*, and filtered sites by minimum and maximum depth across individuals (-min-meanDP 1 and -max-meanDP 264).

To prepare the input file for the principal component analysis (PCA) and admixture analysis, we next filtered by minimum and maximum depth within individuals (-minDP 4 and -maxDP 25) and by missingness across individuals (-max-missing 0.6), and then removed samples with <25% of these sites. To account for the genotype uncertainty inherent in low-coverage data, we generated genotype probabilities using ANGSD 0.911 [6] with the -doPost 1 and -doGeno 32 options. This dataset contained 80,248 variants across 54 *P. maniculatus* samples. For the PCA, we used the ngsCovar function in ngsTools 0.615 [7] to compute a covariance matrix. For the admixture analysis, we generated a beagle file with the -doGlf function in ANGSD, and used the ngsAdmix function in ngsTools to calculate admixture proportions. We plotted PCs and admixture proportions following this tutorial: https://github.com/mfumagalli/ngsTools/blob/master/TUTORIAL.md.

To prepare the input file for the divergence dating analysis, we selected a subset of high coverage samples (3 Saturna Island *P. maniculatus*, 5 Pender Island *P. maniculatus*, 3 mainland *P. maniculatus*, 3 *P. keeni*), and filtered by minimum and maximum depth within individuals (-minDP 5 and -maxDP 25) and missingness across individuals (-max-missing 0.8). This resulted in a dataset of 7,092 variants across 14 samples. We then converted the vcf file to nexus format with custom Python code, and used SNAPP 1.3.0 [8] to create a phylogenetic tree with divergence time estimates. Briefly, we loaded the nexus file into a SNAPP template in BEAUti 2.4.8 [9], assigned species IDs to samples, calculated mutation rates, logged every 500 trees, and used a chain length of 10^6^. We ran 10 separate iterations of the resulting XML file in BEAST 2.4.8 [9], and combined log and tree files across iterations in LogCombiner with a by-iteration burn-in of 10%. We checked for convergence and high ESS values in Tracer 1.6 [10], and used TreeAnnotator to calculate a maximum clade credibility tree with median heights. We produced the final tree using FigTree 1.4.2.

### Establishment of laboratory colonies and animal husbandry

We housed animals on Bed-o’Cobs 1/4” bedding (The Andersons, Maumee, Ohio) in ventilated standard rodent cages (Allentown Inc., Allentown, NJ) on a 16h light: 8h dark cycle at 23°C. Animals were provided with a red translucent polycarbonate hut, Enviro-Dri nesting material, and a cotton nestlet. Animals were given *ad libitum* access to irradiated Prolab Isopro RMH 3000 5P74 (LabDiet) and water. We weaned litters at 23 days of age and kept animals in groups of up to five individuals of the same sex and strain.

### Behavioural experiments

Approximately 3-4 hours before the trial, the female was removed from the resident’s cage, resident and intruder were weighed, and the intruder was marked for easy identification with permanent marker on the tail. At the start of the experiment, we placed the resident’s cage into black blinders, removed the lid and the red hut, and placed a custom-built divider with a gate and a transparent lid onto the cage. The divider set apart a small portion of the cage for the intruder, but was perforated to enable contact. The resident was habituated to the divider for 10 mins. Then we introduced the intruder into the area separated by the divider, and both mice were habituated for another 10 mins. Next, we opened the gate and the intruder was allowed to enter the resident’s area. We video-recorded behaviour from above the cage at 15 fps; trials lasted for 15 mins.

### Statistical analysis

We tested for differences in weight, body composition, and skeletal traits with linear fixed effects models with a strain (island/mainland) by origin (field/lab) interaction. We tested for differences in birth weight of parental strains (or F1 hybrids) using linear fixed effects models with the natural logarithm of weight as the dependent variable and the natural logarithm of litter size and strain (or maternal strain, respectively) as explanatory variables. We tested for differences in growth rate between parental strains (or F1 hybrids) using linear fixed effects models with the natural logarithm of growth rate as the dependent variable and the natural logarithm of litter size and an interaction of strain (or maternal strain, respectively) and day as explanatory variables, and adjusted p-values for multiple testing with the Holm method. Growth rates were obtained by taking the first derivative of cubic smoothing splines fit to the weight data using smoothPspline {pspline}. Our sample size for the F1 hybrids was moderate, therefore we ran additional tests that controlled for family effects. We did not include parent ID (as a proxy for family effects) into the model to avoid model estimation problems introduced by the perfect collinearity of parent ID and strain. Instead, we accounted for family effects by adjusting the standard errors of the model using wild cluster bootstrapping as implemented in cluster.wild.glm {clusterSEs}. To compare litter weight before and after the first milk meal, we used a repeated measures linear mixed effects model with the natural logarithm of weight as dependent variable, the natural logarithm of litter size and an interaction of feeding status (pre-vs. post-feeding) and strain as explanatory variables and litter ID as random effect. We tested for differences in weight by maternal strain in adult hybrid mice with a linear fixed effects model with the natural logarithm of weight as the dependent variable and strain as the explanatory variable. In the cross-fostering experiment, we tested for differences in the natural logarithm of growth rate of conspecific non-fostered and fostered litters with a linear fixed effects model, with the natural logarithm of litter size and an interaction of state (fostered vs. non-fostered) and day as explanatory variables, and adjusted p-values for multiple testing with the Holm method.

To reduce the dimensionality in the behaviour of the wild-caught resident animals, we performed a PCA on behaviours averaged across trials using prcomp {stats}, with variables scaled to unit variance and shifted to be zero centred. We next selected variables that contributed most to the first PC and focused on these behaviours for subsequent analyses. Specifically, we compared levels in these three behaviours (wrestling, chasing, and pindown) in wild-caught residents, captive-born residents one week after they were paired with a female, and breeding pairs after the onset of mating attempts. For this, we used a two-model hurdle approach to test for differences between strains. This included a binary model to test whether different proportions of individuals express a behaviour, and count and duration models to test whether differences exist in the number of bouts and duration once animals express a given behaviour. We initially evaluated the impact of trial number and weight difference between resident and intruder on aggression by comparing repeated measures linear mixed effects models with and without these factors using likelihood ratio tests. Because these factors did not improve models for most behaviours, we averaged data across trials (average number of bouts were rounded) and ran logistic regression models using glm {stats} with binomial (binary) and quasipoisson (count) error distributions and linear fixed effects models (duration). To compare wrestling duration before and after reproduction, we fit a repeated measures mixed effects linear model on the averaged dataset with reproductive status and strain as explanatory variables and resident ID as random effect. To evaluate other potential impacts on the summed duration of wrestling, chasing, and pindown in wild-caught mice, we fit a repeated measures mixed effects linear model with days since capture, days since last litter was sired, weight difference, and strain as explanatory variables and resident ID as random effect.

---

**Suppl. Fig. 1.**
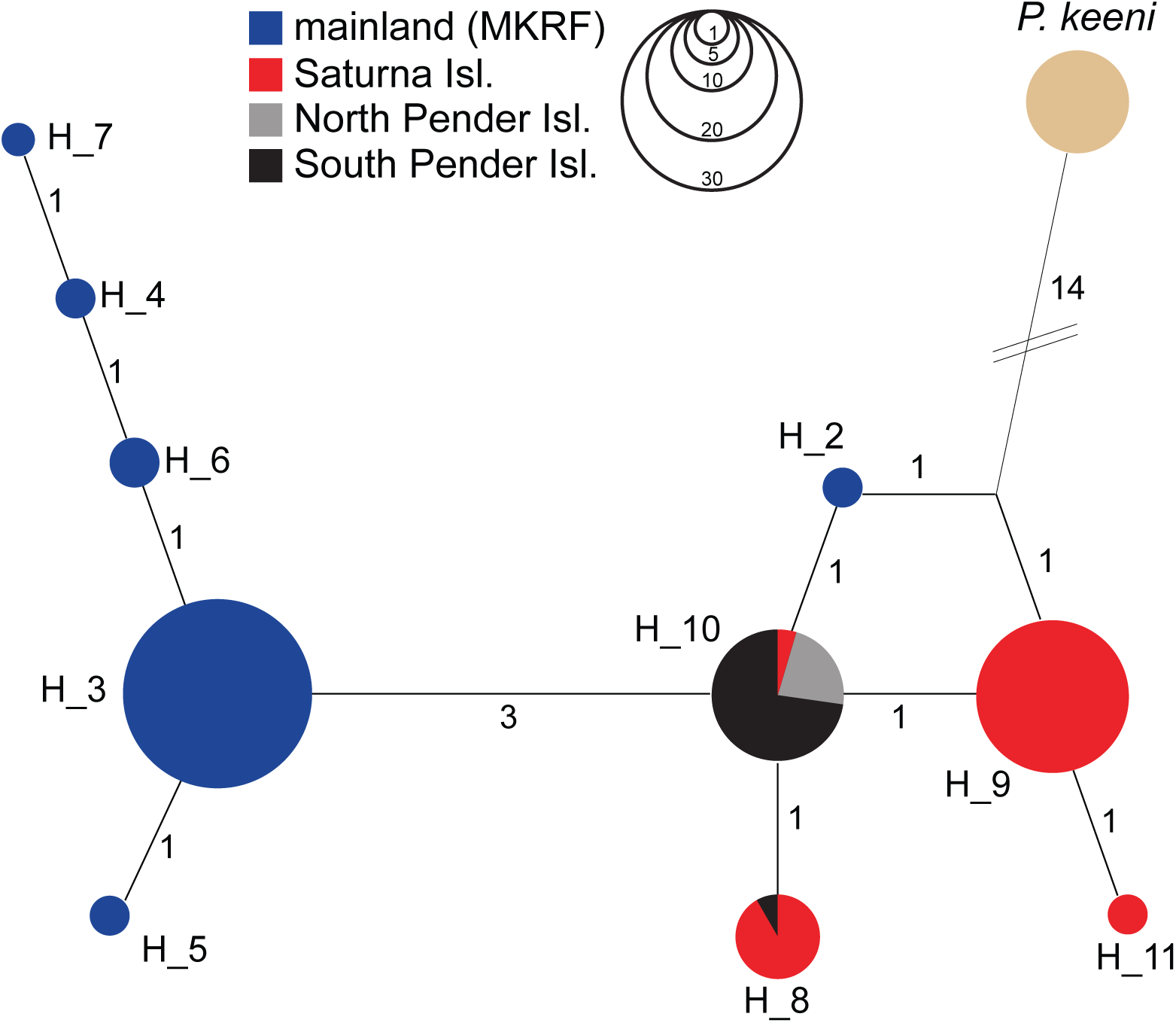
Haplotype network based on 381bp of the mitochondrial cytochrome *b* gene. Numbers on edges indicate the number of DNA substitutions separating adjacent haplotypes. Diameter of nodes is proportional to the number of specimens that carry the respective haplotype (see schematic at the top of the graph).

**Suppl. Fig. 2.**
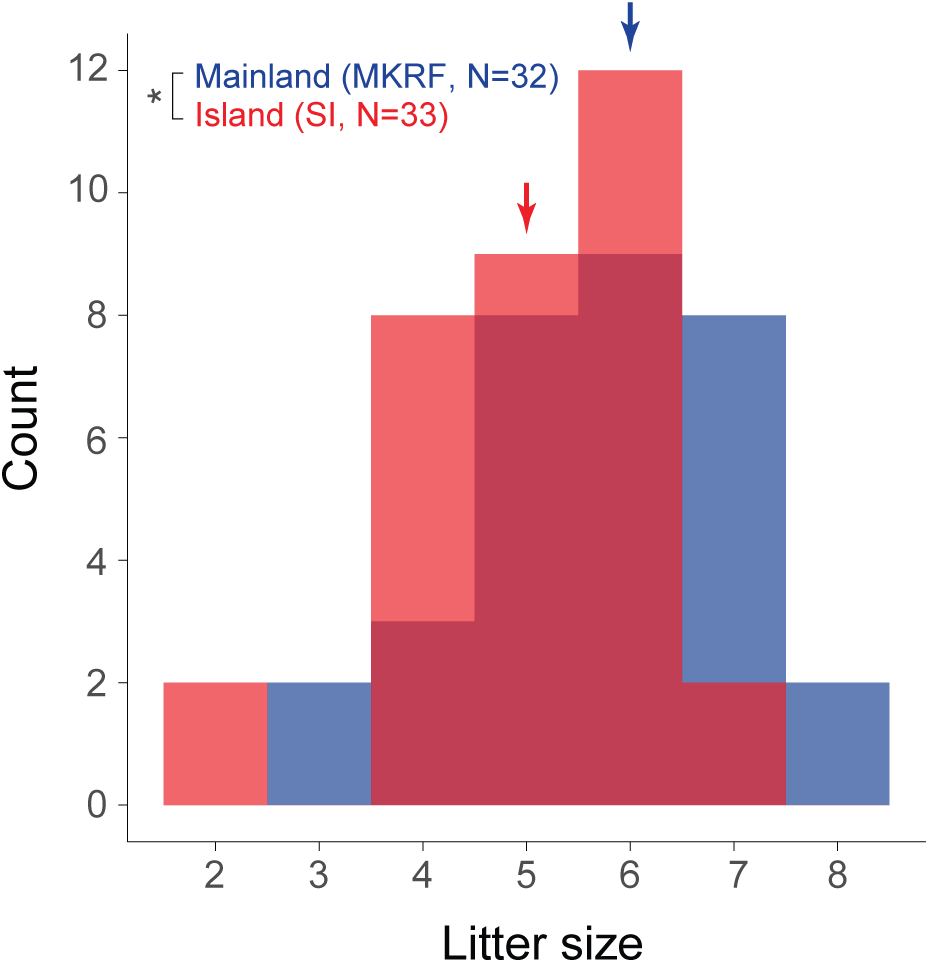
Litter size of island (red) and mainland (blue) mice. Arrows indicate the median litter size for each strain. Statistical significance evaluated by Kruskal-Wallis test. * *P* < 0.05.

**Suppl. Fig. 3.**
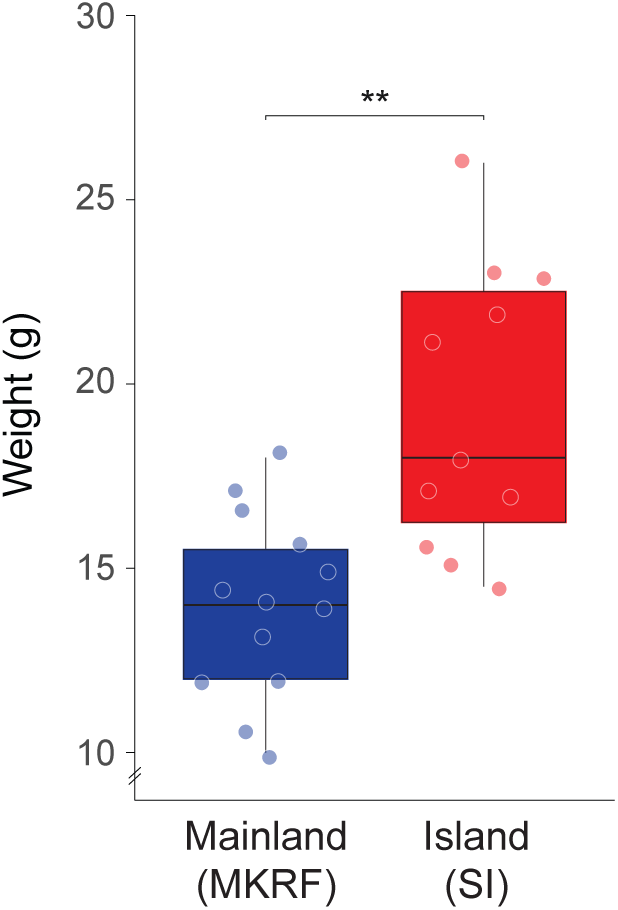
Body weight in adult female wild-caught island (red) and mainland (blue) mice. Statistical significance evaluated by linear fixed effects model (see Methods for details). ** *P* < 0.01.

**Suppl. Fig. 4.**
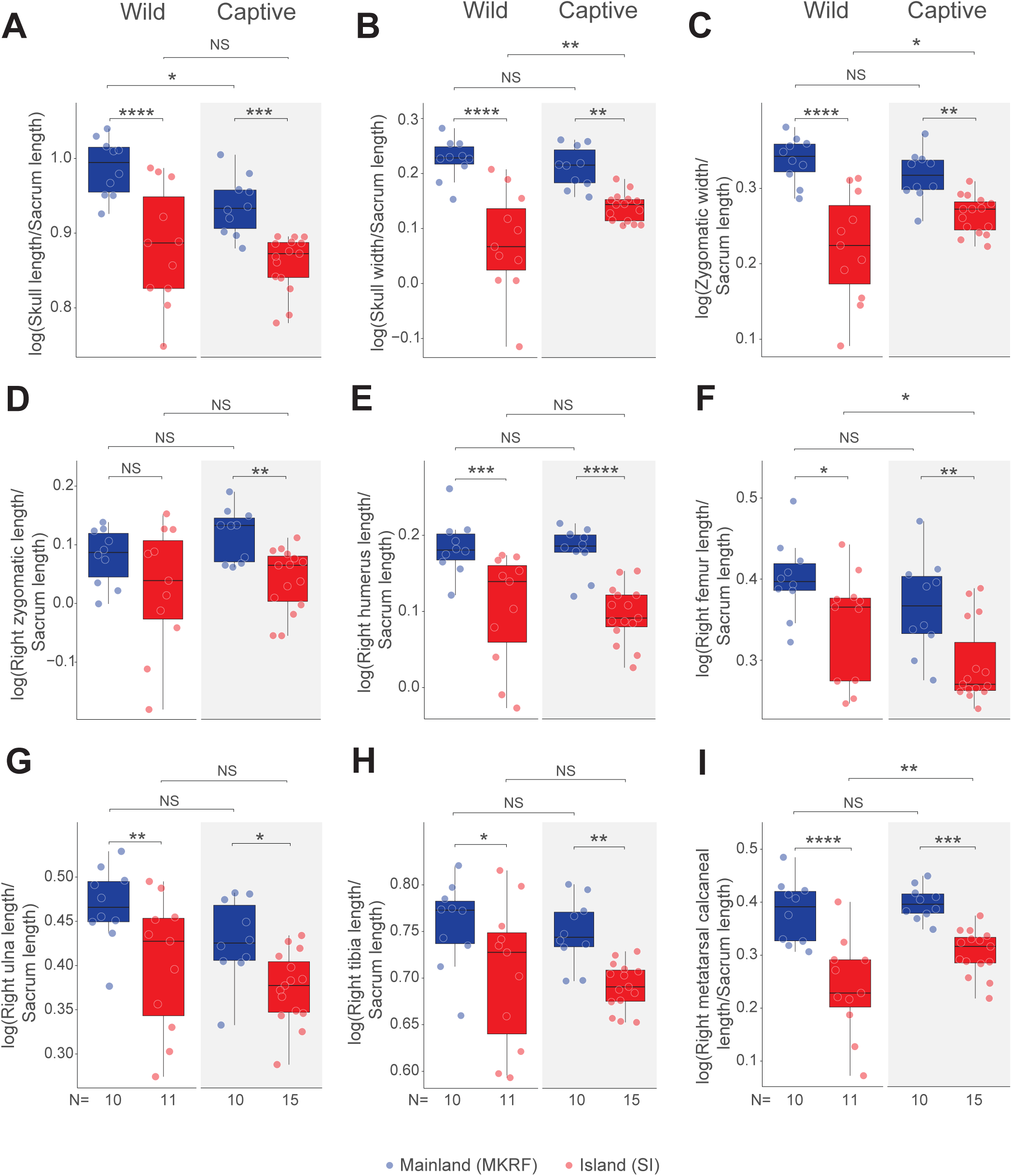
Size-corrected (**A**) skull length, (**B**) skull width, (**C**) zygomatic width, (**D**) right zygomatic length, (**E**) right humerus length, (**F**) right femur length, (**G**) right ulna length, (**H**) right tibia length, and (**I**) right metatarsal calcaneal length in male wild-caught (left) and captive-born (right) island (red) and mainland (blue) mice. Statistical significance evaluated by linear fixed effects model (see Methods for details). NS=not significant, * *P* < 0.05, ** *P* < 0.01, *** *P* < 0.001, **** *P* < 0.0001.

**Suppl. Fig. 5.**
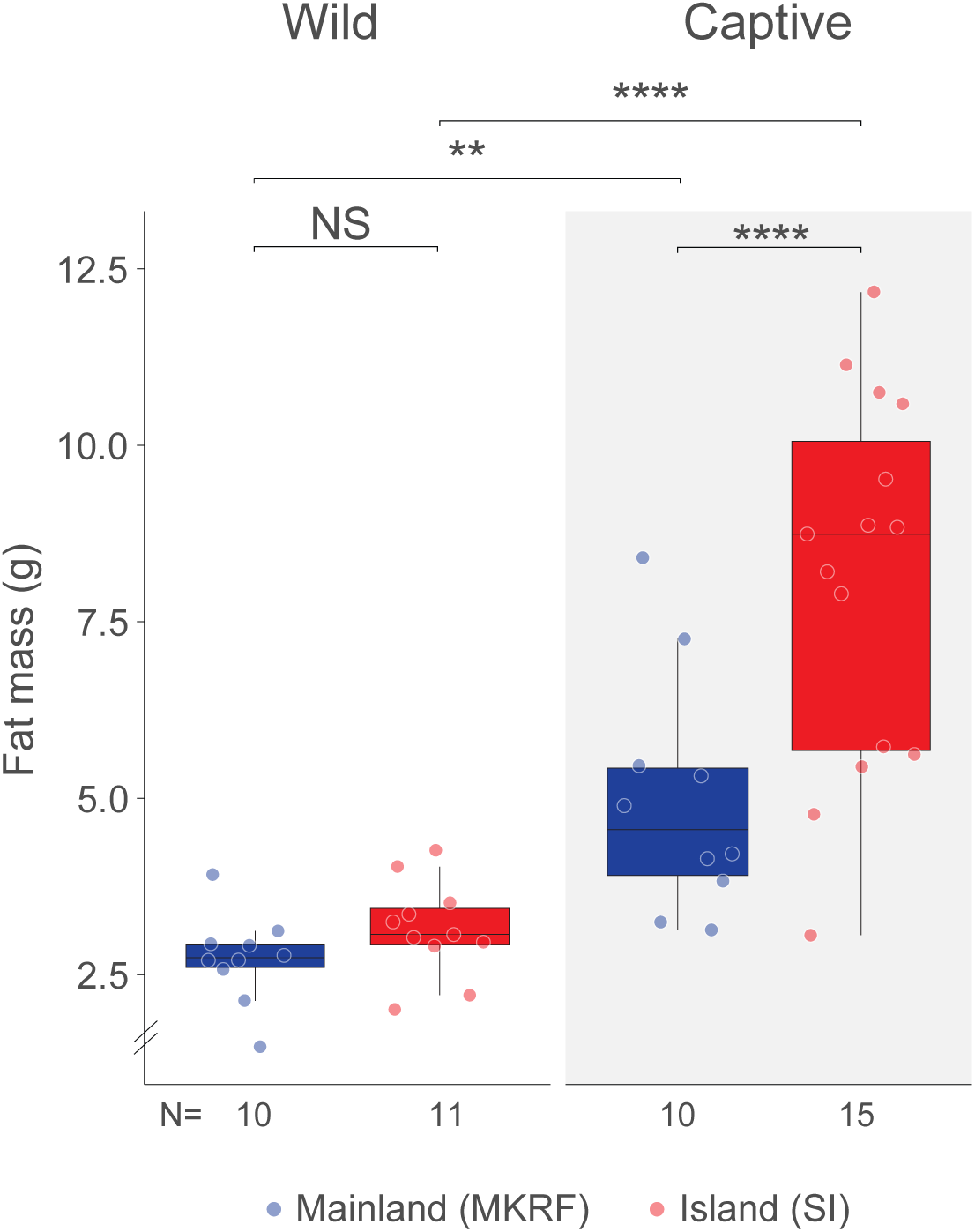
Fat mass in adult male wild-caught (left) and captive-born (right) island (red) and mainland (blue) mice. Statistical significance evaluated by linear fixed effects model (see Methods for details). NS=not significant, ** *P* < 0.01, **** *P* < 0.0001.

**Suppl. Fig. 6.**
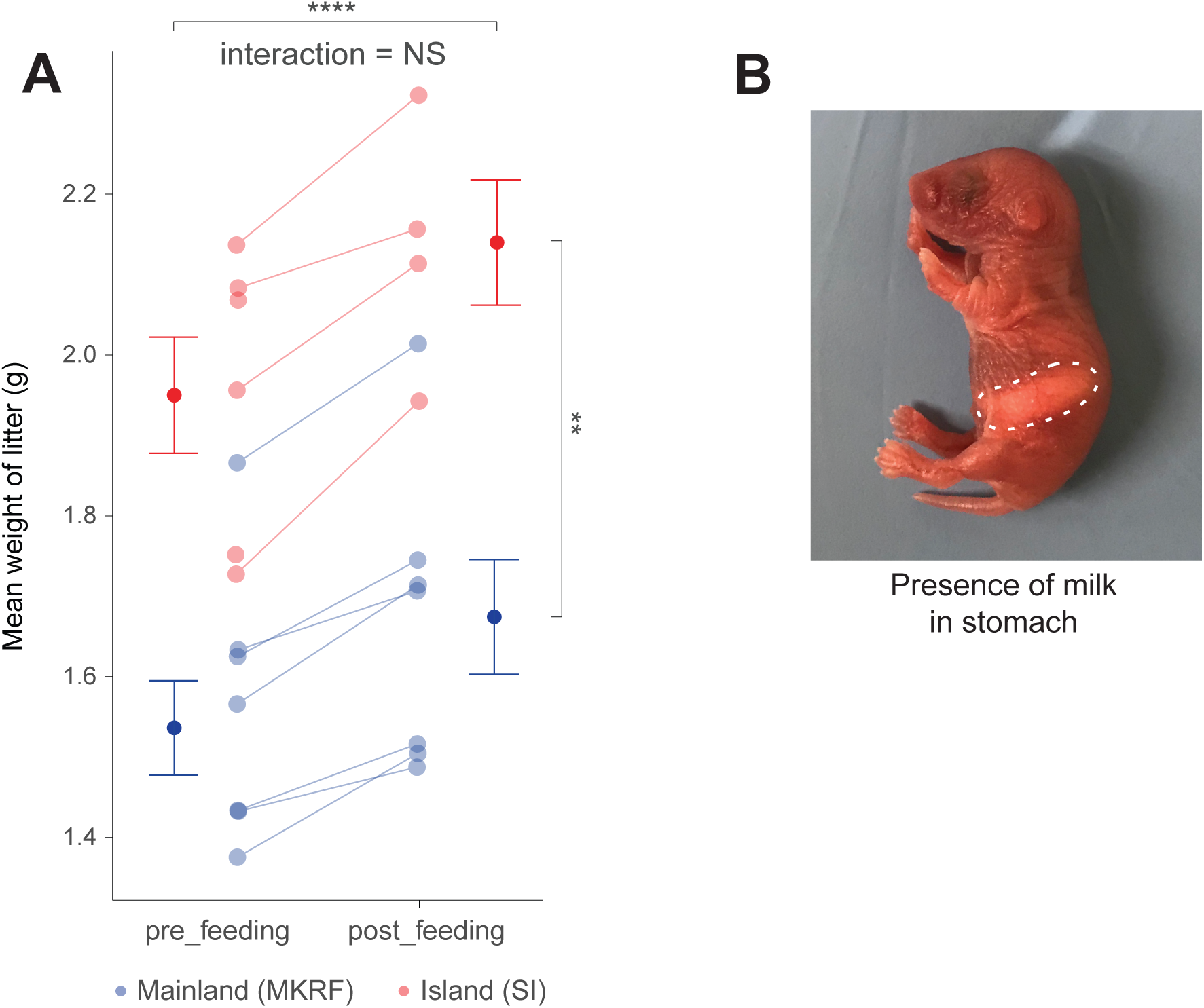
(**A**) Mean weight of island (red) and mainland (blue) litters before and after the first milk meal. Statistical significance evaluated by repeated measures linear mixed effects model (see Methods for details). NS=not significant, ** *P* < 0.01, **** *P* < 0.0001. (**B**) Example of pup with milk present in stomach (outlined).

**Suppl. Fig. 7.**
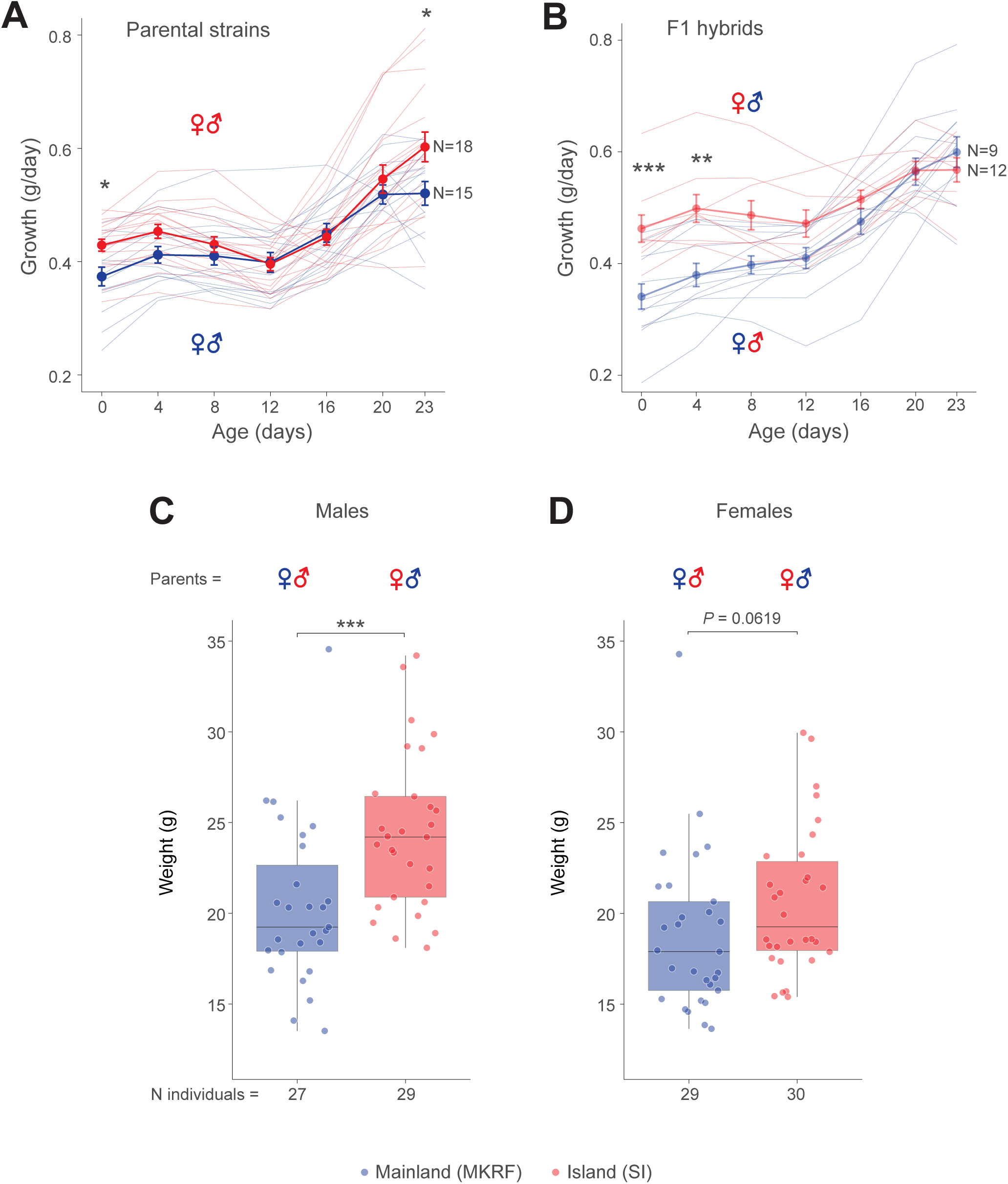
(**A-B**) Growth curves of (**A**) island (red) and mainland (blue) litters and (**B**) F1 hybrid litters by maternal strain. Points and error bars represent mean and SEM of growth of litters. (**C-D**) Weight of adult hybrid mice plotted by maternal strain identity. The natural logarithm of weights of (**C**) males and (**D**) females were separately compared by maternal strain with linear fixed effects models (see Methods for details). * *P* < 0.05, ** *P* < 0.01, *** *P* < 0.001.

**Suppl. Fig. 8.**
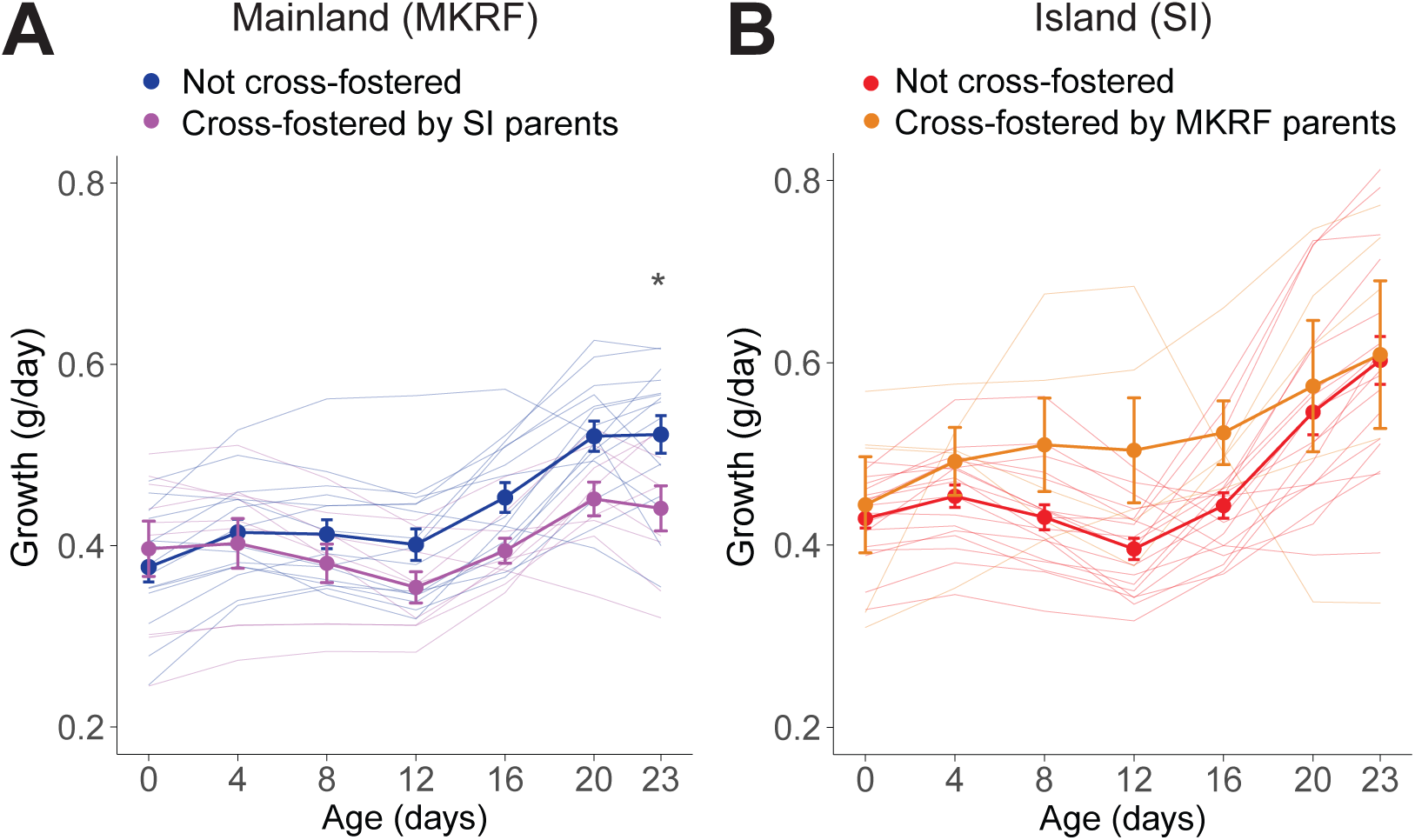
Growth rates of (**A**) mainland litters cross-fostered by island parents (purple), compared to mainland litters that were not cross-fostered (blue), (**B**) island litters cross-fostered by mainland parents (orange), compared to island litters that were not cross-fostered (red). Points and error bars represent mean and SEM of growth of litters. Statistical significance evaluated by linear fixed effects model (see Methods for details). * *P* < 0.05.

**Suppl. Fig. 9.**
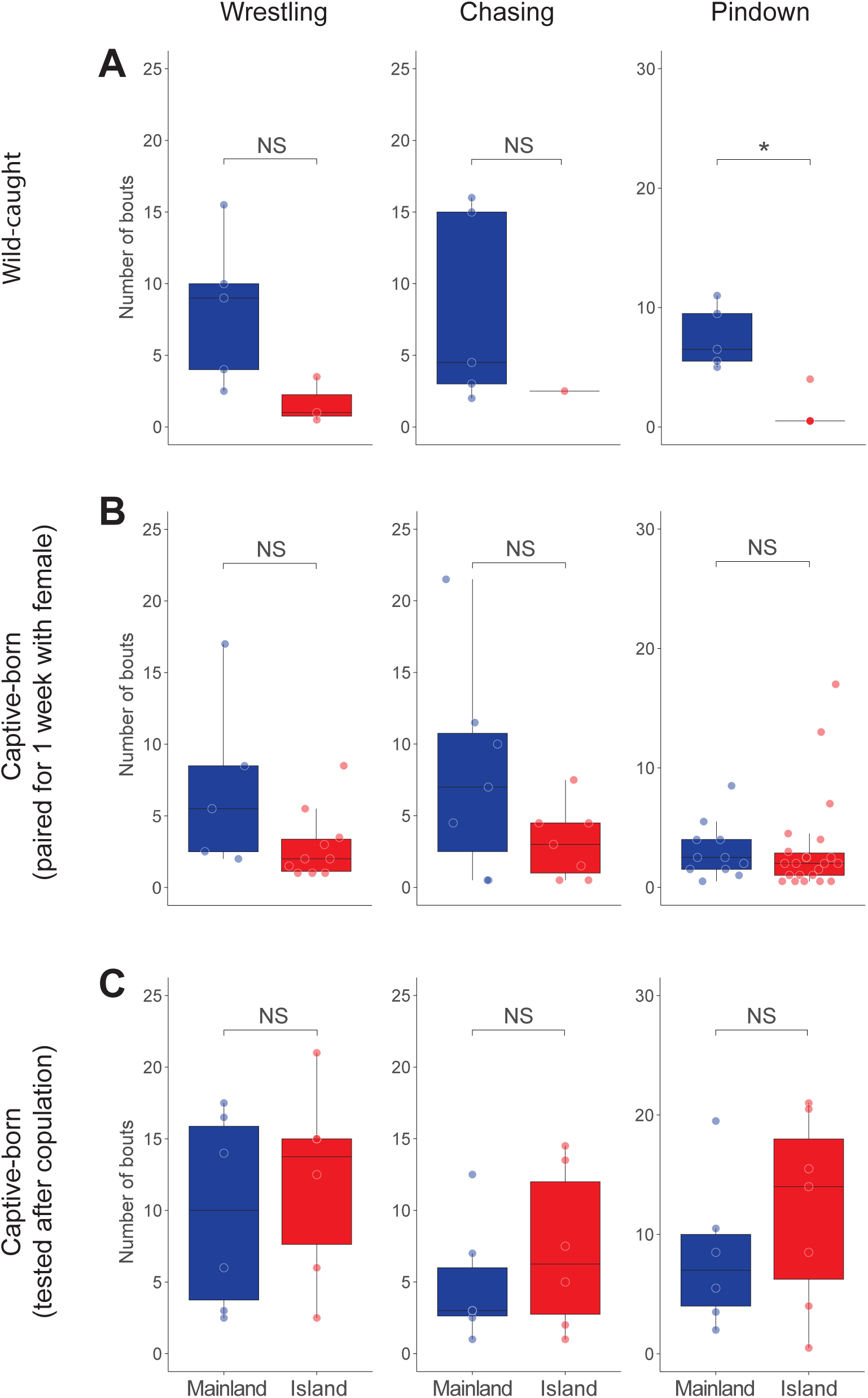
Count data of wrestling, chasing, and pindown behaviours in wild-caught founder animals (**A**), captive-born males paired with a female for one week before testing (**B**), and captive-born breeding males tested after verified copulation (**C**). Statistical significance evaluated by generalized linear models (see Methods for details). NS=not significant, * *P* < 0.05.

**Suppl. Fig. 10.**
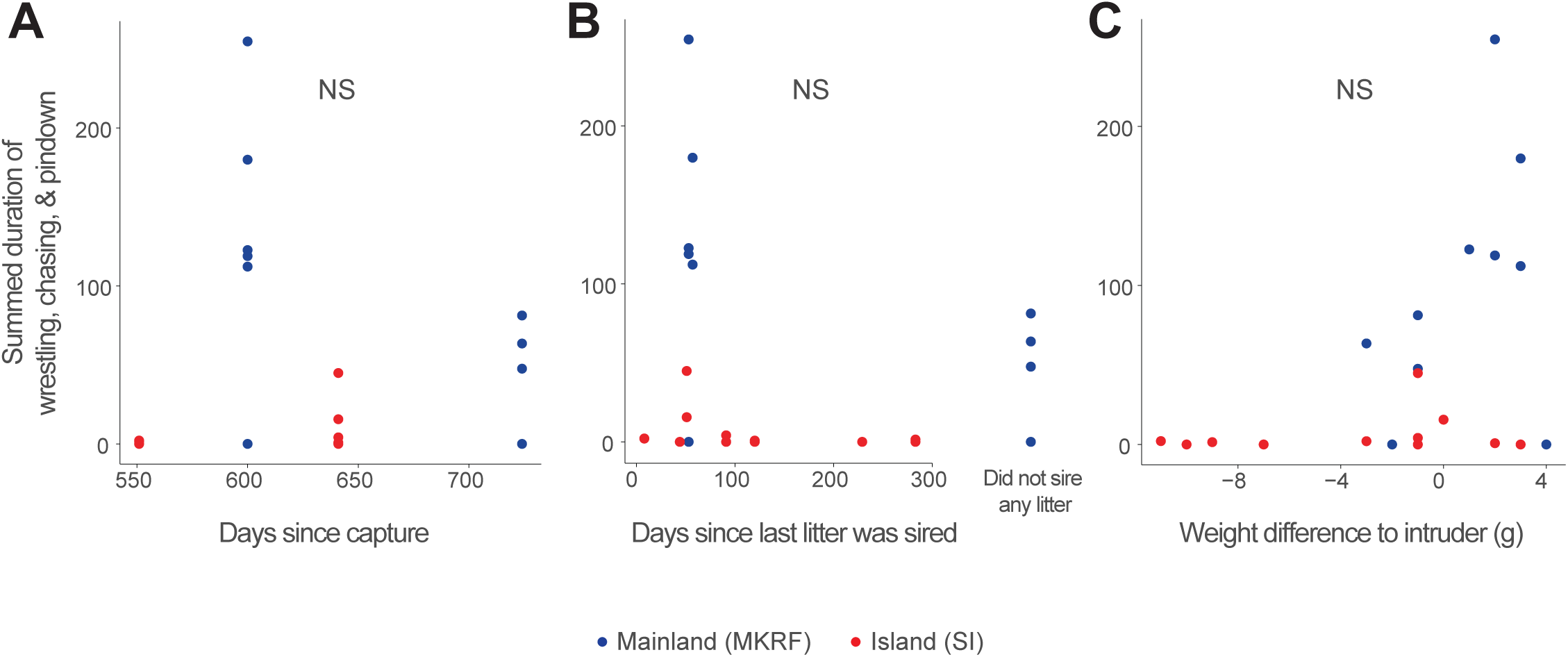
Effect of (**A**) time since capture at testing, (**B**) time since a litter was last sired, and (**C**) weight difference between resident and intruder on the aggressive behaviour of wild-caught deer mice (N=7 island [red], N=6 mainland [blue]). Replicate trials are plotted separately. Statistical significance evaluated with a repeated measures mixed effects linear model (see Methods for details). NS=not significant.

**Suppl. Fig. 11.**
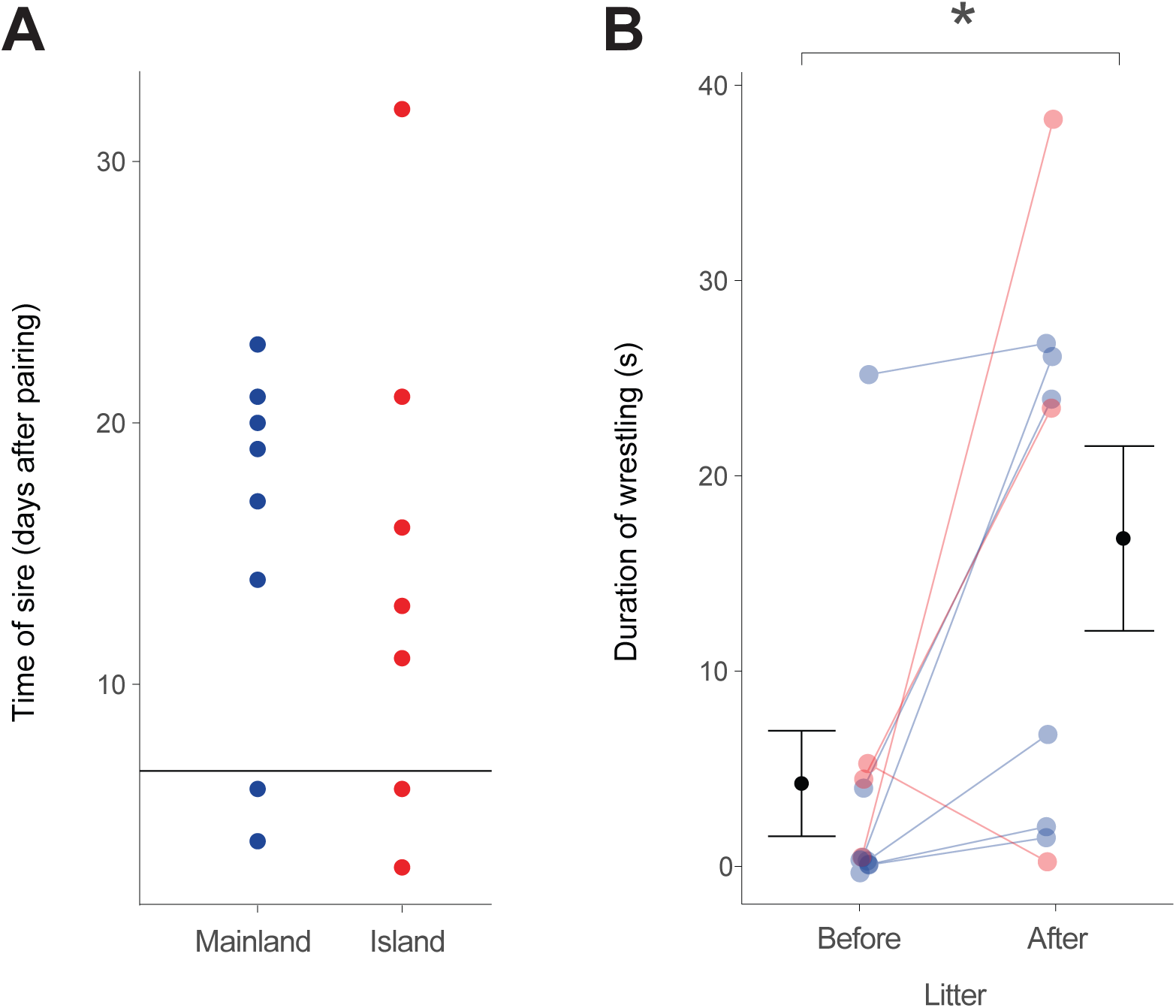
Effect of reproductive experience on territorial aggression. (**A**) Time of mating after pairing in a subset of mice from the experiment in Fig. 5A-B. We backdated for females that went on to give birth when the litter was sired. The horizontal line marks the time when males were tested in the resident-intruder assay (7 days after pairing); at this time, few males in this subset had sired litters. (**B**) We re-tested a subset of these males after they had sired a litter, and compared wrestling duration before and after siring a litter with a repeated measures linear mixed effects model (see Methods for details). Only mice were included that had not sired a litter by the time of first testing. Mean and SEM across strains is shown in black. * *P* < 0.05.

**Suppl. Fig. 12.**
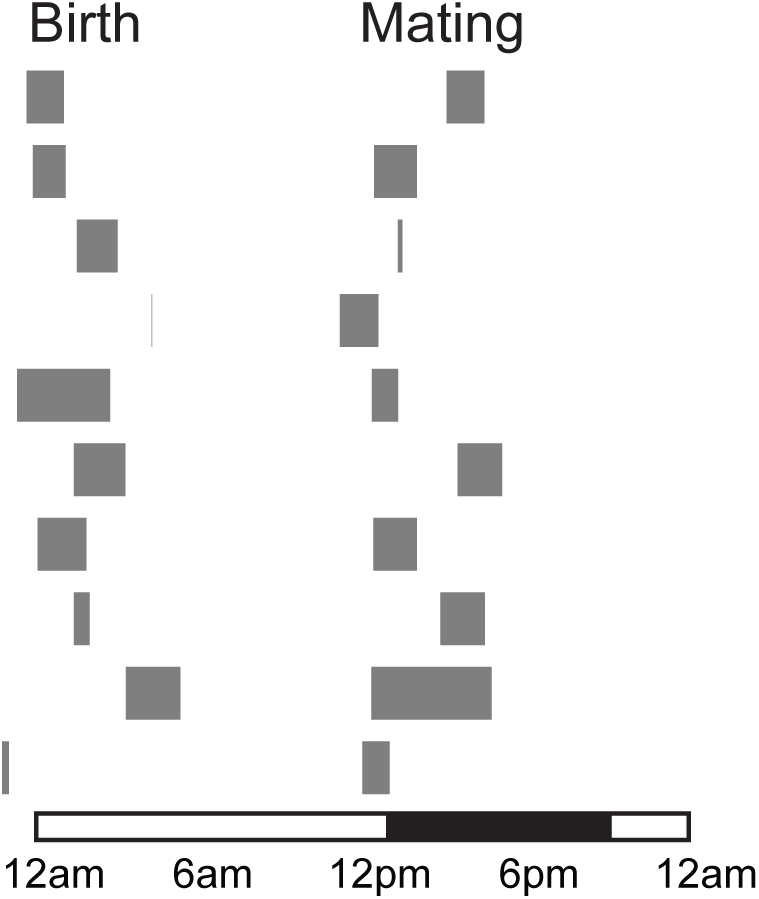
Schematic showing birth and mating events in breeding pairs. Each row corresponds to one pair. The grey boxes indicate the maximum duration of behaviours (it was not possible to observe females continuously, e.g. when they were inside the nest at the time of birth). The light cycle (16h:8h, light:dark) is shown at the bottom.

